## Supplemental Article S1 for "The energetic effect of hip flexion and retraction in walking at different speeds: a modeling study"

### A. Equations of motion

Based on the framework of Casius et al. (2004), the equations of motion at four different phases were derived by solving the unknown angular accelerations $\ddot{\phi}_i$ and the unknown forces $F_{ix}, F_{iy}$ (either ground reaction force or joint reaction force) acting on each segment, with input the external torques of ankle and hip joint. To solve these 12 unknowns, the 12 equations of motion were derived using Newton's second law regarding the linear accelerations (horizontal and vertical direction) and using Euler's equation regarding angular acceleration for each segment. The adjacent segments interact with each other by the coupling of (opposite) joint reaction forces and net joint torques. The equations of motion for the toe-constrained phase are listed in Eq. A1 as an example.

$$
\begin{aligned}
&F_{1x} - F_{2x} = m_1\ddot{x}_1\\
&F_{2x} - F_{3x} = m_2\ddot{x}_2\\
&F_{3x} - F_{4x} = m_3\ddot{x}_3\\
&F_{4x} = m_4\ddot{x}_4\\
&F_{1y} - F_{2y} + m_1 g = m_1\ddot{y}_1\\
&F_{2y} - F_{3y} + m_2 g = m_2\ddot{y}_2\\
&F_{3y} - F_{4y} + m_3 g = m_3\ddot{y}_3\\
&F_{4y} + m_4 g = m_4\ddot{y}_4\\
&F_{1x}d_1\sin\phi_1 - F_{1y}d_1\cos\phi_1 + F_{2x}(l_1 - d_1)\sin\phi_1 - F_{2y}(l_1 - d_1)\cos\phi_1 - \tau_1^a\\
&= j_1\ddot{\phi}_1\\
&F_{2x}d_2\sin\phi_2 - F_{2y}d_2\cos\phi_2 + F_{3x}(l_2 - d_2)\sin\phi_2 - F_{3y}(l_2 - d_2)\cos\phi_2 + \tau_1^a - \tau_2^h\\
&= j_2\ddot{\phi}_2\\
&F_{3x}d_3\sin\phi_3 - F_{3y}d_3\cos\phi_3 + F_{4x}(l_3 - d_3)\sin\phi_3 - F_{4y}(l_3 - d_3)\cos\phi_3 + \tau_2^h - \tau_3^a\\
&= j_3\ddot{\phi}_3\\
&F_{4x}d_4\sin\phi_4 - F_{4y}d_4\cos\phi_4 + \tau_3^a = j_4\ddot{\phi}_4.
\end{aligned}
\tag{A1}
$$

To transform the linear acceleration terms $\ddot{x}_i$, $\ddot{y}_i$ also to functions of these unknowns, we used the simple algebraic relations $x_1 = d_1\cos\phi_1,\ldots,\ x_4 = l_1\cos\phi_1 + l_2\cos\phi_2 + l_3\cos\phi_3 + d_1\cos\phi_1$ and $y_1 = d_1\sin\phi_1,\ldots, y_4 = l_1\sin\phi_1 + l_2\sin\phi_2 + l_3\sin\phi_3 + d_4\sin\phi_4$, and their double derivatives. The original 12 equations can now be derived in matrix form

$A \cdot x = b,$

where $A$ is given by

$$\begin{bmatrix}
1 & -1 & 0 & 0 & 0 & 0 & 0 & 0 & m_1 d_1 \sin\phi_1 & 0 & 0 & 0 \\
0 & 1 & -1 & 0 & 0 & 0 & 0 & 0 & m_2 l_1 \sin\phi_1 & m_2 d_2 \sin\phi_2 & 0 & 0 \\
0 & 0 & 1 & -1 & 0 & 0 & 0 & 0 & m_3 l_1 \sin\phi_1 & m_3 l_2 \sin\phi_2 & m_3 d_3 \sin\phi_3 & 0 \\
0 & 0 & 0 & 1 & 0 & 0 & 0 & 0 & m_4 l_1 \sin\phi_1 & m_4 l_2 \sin\phi_2 & m_4 l_3 \sin\phi_3 & m_4 d_4 \sin\phi_4 \\
0 & 0 & 0 & 0 & 1 & -1 & 0 & 0 & -m_1 d_1 \cos\phi_1 & 0 & 0 & 0 \\
0 & 0 & 0 & 0 & 0 & 1 & -1 & 0 & -m_2 l_1 \cos\phi_1 & -m_2 d_2 \cos\phi_2 & 0 & 0 \\
0 & 0 & 0 & 0 & 0 & 0 & 1 & -1 & -m_3 l_1 \cos\phi_1 & -m_3 l_2 \cos\phi_2 & -m_3 d_3 \cos\phi_3 & 0 \\
0 & 0 & 0 & 0 & 0 & 0 & 0 & 1 & -m_4 l_1 \cos\phi_1 & -m_4 l_2 \cos\phi_2 & -m_4 l_3 \cos\phi_3 & -m_4 d_4 \cos\phi_4 \\
d_1 \sin\phi_1 & (l_1 - d_1)\sin\phi_1 & 0 & 0 & -d_1 \cos\phi_1 & -(l_1 - d_1)\cos\phi_1 & 0 & 0 & -j_1 & 0 & 0 & 0 \\
0 & d_2 \sin\phi_2 & (l_2 - d_2)\sin\phi_2 & 0 & 0 & -d_2 \cos\phi_2 & -(l_2 - d_2)\cos\phi_2 & 0 & 0 & -j_2 & 0 & 0 \\
0 & 0 & d_3 \sin\phi_3 & (l_3 - d_3)\sin\phi_3 & 0 & 0 & -d_3 \cos\phi_3 & -(l_3 - d_3)\cos\phi_3 & 0 & 0 & -j_3 & 0 \\
0 & 0 & 0 & d_4 \sin\phi_4 & 0 & 0 & 0 & -d_4 \cos\phi_4 & 0 & 0 & 0 & -j_4
\end{bmatrix}$$

and b is given by

$$\begin{bmatrix}
-m_1 d_1 \dot{\phi}_1^2 \cos\phi_1 \\
-m_2 l_1 \dot{\phi}_1^2 \cos\phi_1 - m_2 d_2 \dot{\phi}_2^2 \cos\phi_2 \\
-m_3 l_1 \dot{\phi}_1^2 \cos\phi_1 - m_3 l_2 \dot{\phi}_2^2 \cos\phi_2 - m_3 d_3 \dot{\phi}_3^2 \cos\phi_3 \\
-m_4 l_1 \dot{\phi}_1^2 \cos\phi_1 - m_4 l_2 \dot{\phi}_2^2 \cos\phi_2 - m_4 l_3 \dot{\phi}_3^2 \cos\phi_3 - m_4 d_4 \dot{\phi}_4^2 \cos\phi_4 \\
-m_1 d_1 \dot{\phi}_1^2 \sin\phi_1 - m_1 g \\
-m_1 l_1 \dot{\phi}_1^2 \sin\phi_1 - m_2 d_2 \dot{\phi}_2^2 \sin\phi_2 - m_2 g \\
-m_1 l_1 \dot{\phi}_1^2 \sin\phi_1 - m_2 l_2 \dot{\phi}_2^2 \sin\phi_2 - m_3 d_3 \dot{\phi}_3^2 \sin\phi_3 - m_3 g \\
-m_1 l_1 \dot{\phi}_1^2 \sin\phi_1 - m_2 l_2 \dot{\phi}_2^2 \sin\phi_2 - m_3 l_3 \dot{\phi}_3^2 \sin\phi_3 - m_4 d_4 \dot{\phi}_4^2 \sin\phi_4 - m_4 g \\
\tau_1^a \\
\tau_2^h - \tau_1^a \\
\tau_3^a - \tau_2^h \\
-\tau_3^a
\end{bmatrix}.$$

The equations of motion for other phases can be derived similarly. For the heel-constrained phase and the foot-constrained phase, the only difference is that the ground reaction force $F_{1x}, F_{1y}$ is not acting on the most proximal end (stance toe). Thus, we shifted the ground reaction forces acting on the heel (or any point on the foot) to the toe and added an 'external' torque on the stance foot to equalize the ground reaction force on the heel. This new unknown 'external' torque can subsequently be solved from the magnitude of the ground reaction forces and its moment arm. For the foot-constrained phase, the additional constraint is that the angular acceleration of the stance foot is zero. For the complete equations of motion, see our shared code https://drive.google.com/drive/folders/1Xduqtn1F-TEypfJ6ifbv6u6hMLbCLVky?usp=sharing.

The additional unknown in double stance phase can be solved by constraining the hind toe to a fixed position. To describe the collision equations of heel strike and toe-strike, we applied the law of conservation of angular momentum about the (new) stance foot before and after heel strike. We used the same modeling framework as in the single stance phase, but took the infinitesimal time integral of the coupled differential equations. As a result, the unknowns became (ground-reaction and joint-reaction) impulses and the angular velocity changes for all segments. Since the collision was modelled to be instantaneous, all impulses of finite forces and torques (e.g. gravity and actuator torques) and all angular position changes $[\Delta\phi_1\ \Delta\phi_2\ \Delta\phi_3\ \Delta\phi_4]^T$ were set to zero.

### B. Ground impulsive work

A simplified analytical model for computing the impulse work (or the opposite of collision loss) is illustrated in Figure S1, where we focus on the transition of leading leg states at heel strike and all other segments are neglected. The more complicated four-segment case will be analyzed at the end of this section. The leading leg has center-of-mass $M + m$ located at $r$ distance from the heel, with pre-collision angular velocity $\dot{\varphi}$, pre-collision velocity $v_{c-}$ and post-collision velocity $v_{c+}$. For this simplified single segment model, the (ground) impulsive work at heel strike is given by the difference of kinetic (including linear and rotational) energy before and after heel strike

$$W_{\text{impulsive}} = \frac{1}{2}(M + m)(v_{c+}^2 - v_{c-}^2) + \frac{1}{2}J(\varphi_+^2 - \varphi_-^2), \tag{B1}$$

where $J$ is moment of inertia relative to the center-of-mass $M + m$. The horizontal and vertical ground reaction impulse at the heel is given by

$$\begin{aligned} S_x &= (M + m)(v_{cx+} - v_{cx-}), \\ S_y &= (M + m)\left(v_{cy+} - v_{cy-}\right). \end{aligned} \tag{B2}$$

The ground reaction impulse $S_{\text{GRF}}$ changes the angular velocity of the segment instantaneously, indicated by the angular impulse given by

$$S_\varphi = J(\dot{\varphi}_+ - \dot{\varphi}_-) = r \sin\varphi\, S_x - r\cos\varphi\, S_y. \tag{B3}$$

Since the ground reaction impulse $S_{\text{GRF}}$ is the only external impulse on the segment, the change of CoM linear momentum in the direction perpendicular to the segment (see Fig. S1) is given

$$\frac{S_\varphi}{r} = \frac{J(\dot{\varphi}_+ - \dot{\varphi}_-)}{r} = \sin\varphi\, S_x - \cos\varphi\, S_y. \tag{B4}$$

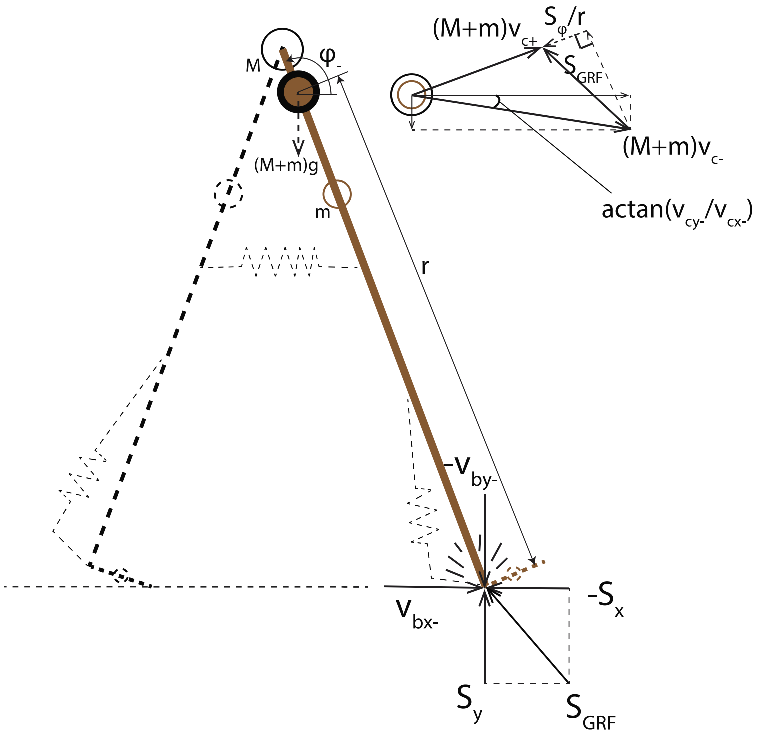


**Fig. S1.** The simplified one-segment model of the relation between the ground reaction impulse and the CoM momentum.

The impulsive work can now be calculated as

$$W_{\text{impulsive}} = \frac{1}{2}(M+m)(v_{cx+}^2 - v_{cx-}^2) + \frac{1}{2}(M+m)\left(v_{cy+}^2 - v_{cy-}^2\right) + \frac{1}{2}J(\dot{\varphi}_+^2 - \dot{\varphi}_-^2)$$
$$= \frac{1}{2}S_x(v_{cx+} + v_{cx-}) + \frac{1}{2}S_y\left(v_{cy+} + v_{cy-}\right) + \frac{1}{2}S_\varphi(\dot{\varphi}_+ + \dot{\varphi}_-) = S_x\bar{v}_{cx} + S_y\bar{v}_{cy} + S_\varphi\bar{\dot{\varphi}}, \quad \text{(B5)}$$

where the bar indicates the averaged states before and after heel strike. The horizontal and vertical velocity of the base of the segment (i.e., heel) is given

$$\bar{v}_{bx} = \frac{1}{2}v_{bx-} = \frac{1}{2}(v_{cx-} + \dot{\varphi}_- r\sin\varphi_-),$$
$$\bar{v}_{by} = \frac{1}{2}v_{by-} = \frac{1}{2}\left(v_{cy-} - \dot{\varphi}_- r\cos\varphi_-\right). \quad \text{(B6)}$$

Based on the equations (B4-B6), the impulsive work can be derived as

$$W_{\text{impulsive}} = \frac{1}{2}S_x v_{bx-} + \frac{1}{2}S_y v_{by-}, \quad \text{(B7)}$$

where the impulsive work is half the product of impact velocity and ground reaction impulse.

Note that Eq. B7 to compute the impulsive work is based on the simplified model of a single segment. When considering the complete four segment model, there are also ground reaction impulses at the trailing toe and joint reaction impulses at the hip during heel strike. Nevertheless, there is no external work done on the whole system from these additional impulses, as there is no velocity change at the toe. Therefore, Eq. B7 to compute the impulsive work also holds for the four-segment system in our model, or any multi-segment model; see Font-Llagunes & Kövecses (2009).

### C. Mechanical work analysis

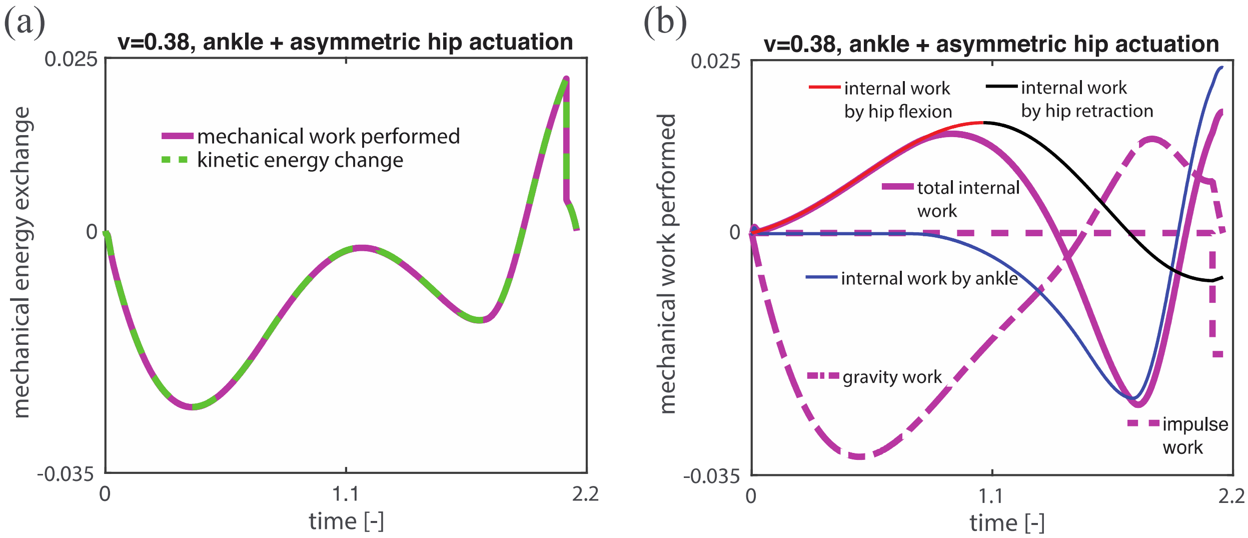


**Fig. S2.** (a) The total mechanical work performed over a gait cycle equals the kinetic energy change; (b) the total mechanical work performed over a gait cycle consists of the mechanical work performed by gravity, (ground) impulsive work, and internal work from ankle, hip flexion and retraction actuation. The periodic gait is an ankle, hip flexion and retraction actuated gait at a speed of 0.38.

### D. MCOT for free symmetric hip actuation cost

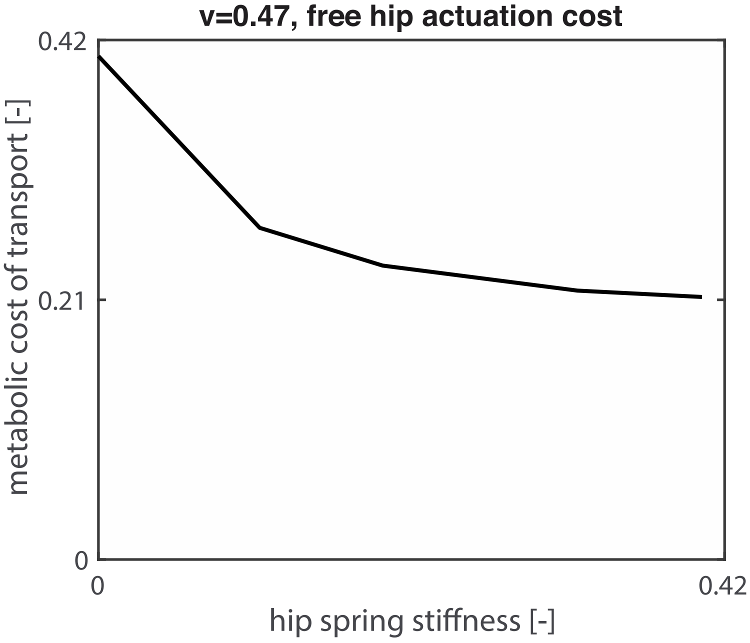


**Fig. S3.** The effects of hip spring stiffness on the MCOT at a high speed 0.47, assuming that hip actuation cost is free. Increasing hip spring stiffness leads to a monotonically decreasing MCOT.

### References

Casius, R., Bobbert, M., & van Soest, K. J. (2004). Forward Dynamics of Two-Dimensional Skeletal Models. A Newton-Euler Approach. *Journal of Applied Biomechanics*, *20*, 421–449. https://doi.org/10.1123/jab.20.4.421

Font-Llagunes, J. M., & Kövecses, J. (2009). Dynamics and energetics of a class of bipedal walking systems. *Mechanism and Machine Theory*, *44*(11). https://doi.org/10.1016/j.mechmachtheory.2009.05.003
